# Population decoding of sound source location by receptive field neurons in the mouse superior colliculus

**DOI:** 10.64898/2026.01.26.701861

**Authors:** Brian R. Mullen, Alan M. Litke, David A. Feldheim

## Abstract

Identifying the location of a sound source in a complex environment and assessing its importance can be crucial for survival. The superior colliculus (SC), a midbrain structure involved in sensorimotor functions, contributes to sound localization and contains auditory responsive neurons that have spatially restricted receptive fields (RFs) that are organized into a topographic map along the azimuth. However, individual auditory SC neurons have large spatial RFs, are noisy, and do not respond to the same stimulus at each trial. Therefore, when an animal is presented with a “single trial” sound, and it needs to rely on a *single* neuron to locate the sound source direction, the location measurement may be erroneous, missing, or have poor spatial resolution. It is expected that a more reliable and accurate determination of the sound source location will come from a population of neurons. We therefore built a population pattern Maximum Likelihood Estimation (MLE) decoder to build a model that can accurately predict the location of a stimulus given the population response. We compared three models that use either strict firing rate (FR), weighting based on equal (EW) or mutual information (MIW) and show that the MIW model works best, needing only 92 neurons to localize a stimulus with behaviorally relevant precision. Furthermore, by comparing the model’s fit using the responses from non-RF and RF auditory neurons, we show that only RF neurons contain the information needed to localize a sound source. These results are consistent with the hypothesis that the SC uses a population of RF neurons to determine sound source location.

**Author Summary:** Being able to tell where a sound is coming from and how important it is can be critical for survival. The superior colliculus, a midbrain region involved in orienting behaviors, contains neurons that respond best to sounds coming from specific locations. This suggests that the combined activity of many neurons in the SC is used to determine sound location from a single sound event. To test this idea, we modeled responses from mouse SC neurons while sounds were played from different positions in space, both along the elevation and horizon. A model that weighted the most informative neurons performed best in both directions needing only 92 neurons to localize a stimulus with behaviorally relevant precision along the azimuth. Comparing the model’s fit using the responses from non-RF and RF auditory neurons, we show that only RF neurons contain the information needed to localize a sound source Overall, our findings show that the SC can accurately locate sounds in both horizontal and vertical space using a population-based strategy, providing a simple and effective solution for rapid sound localization.

## Introduction

The ability to locate the source of a sound is critical for the survival of a wide range of species. In nocturnal animals, such as mice, it provides essential information about the location of both predators and prey. In visual mammals, such as humans, maps of auditory space integrate with visual maps to improve the sensitivity of detecting stimuli and solve ambiguities that arise from processing just a single attribute (Stein and Alex Meredith, 1993). Incoming sound waves are modulated by the head and pinna before activating the cochlea, which contains frequency-tuned neurons that are arranged in a tonotopic order. This organization is maintained as sound information is processed in the inferior colliculus (IC), medial geniculate nucleus, and primary auditory cortex (A1), resulting in neurons that are tuned to frequency but generally contain broad spatial receptive fields (RFs) that encompass a large area of contralateral or ipsilateral space (van der Heijden et al., 2019). Despite this lack of spatial tuning, it has been argued that sound location can be computed by a population rate-code strategy that compares levels of neural activity between the opposing two channels that vary across sound space, whereby the highest firing rate difference is located in the periphery where interaural time and level differences are greatest (Derey et al., 2016; Groh et al., 2003; Lesica et al., 2010; McAlpine et al., 2001; Ortiz-Rios et al., 2017).

The superior colliculus (SC), on the other hand, has been shown to contain auditory neurons with spatially restricted RFs (Ito et al., 2020; Palmer and King, 1982; Withington-Wray et al., 1990). We recently performed an extensive physiological analysis of SC neurons in response to broadband (5-80 kHz) sound coming from different virtual locations in awake behaving mice (Ito et al., 2020). We determined that 21.4% of auditory responsive neurons in the mouse SC have spatially restricted RFs, and that these RFs are distributed along the SC such that they form a topographic map of azimuthal, but not elevation, space. The existence of a topographic map suggests that a population code, instead of or in addition to a rate code, could be employed to localize a sound source when presented with a single trial (Day and Delgutte, 2013). Therefore, we built and compared population coding models to determine how well they can localize sound from a single trial with locations that vary along both azimuth and elevation. Specifically, we used a population pattern maximum likelihood estimate (MLE) decoder model to predict the location of a stimulus given the population response.

We compared three models that sum each of the RF neuron’s sound source likelihood weighted by the strict firing rate (FR), equalized weight (EW), or mutual information (MIW). We find that the MIW model is best at predicting the azimuthal and elevation location of a stimulus given the responses, achieving accuracy similar to those estimated from mouse sound discrimination assays (Behrens and Klump, 2016). Furthermore, by comparing the model’s fit using the responses from neurons with spatially restricted RFs (RF neurons) and those without spatially restricted RFs (non-RF neurons), we show that only RF neurons contain the information needed to localize a sound source. Finally, we bootstrap the RF neuronal population to show that as few as 92 neurons can reliably predict the azimuthal location of an object with an 18° azimuthal error, whereas 283 neurons predict the elevation location with the same angular error. Taken together, these results suggest that the population response of a limited number of auditory responsive RF neurons in the SC can be used to perform behaviorally relevant sound localization computations from a single sound source presentation.

## Results

We concatenated two previously described datasets to build and test different population decoding models using SC neurons that have spatially restricted RFs and represent all anatomical and auditory spatial positions (Ito et al., 2021, 2020). Each dataset had 30 trials of 100 ms burst of white noise (5-80 kHz) for each of 85 equally spaced positions (17 azimuthal x 5 elevation positions). For model testing, we determined which neurons had spatially restricted receptive fields (hereafter called RF neurons) that originated in the SC but did not have axonal wave forms. Our analysis presented here is based on the responses of RF neurons to 20 of the 30 trials at each of the 85 positions, and we set aside the remaining 10 trials to test the models. We also limited our analysis to the first 20 ms after the response (see methods).

To demonstrate that the neurons used for our modeling were diverse with respect to their location, we mapped the physical location of all auditory responsive neurons from 22 experiments (seventeen single insertion experiments and five triple [medial, central, and lateral] insertion experiments) onto a single SC (Figure 1A, B). This revealed that the majority of auditory responsive neurons are located in the deep SC (Fig. 1A), and are distributed along the A-P axis in the central SC, with some neurons located in the medial and lateral SC (Fig. 1B). To determine the areas of auditory space to which these neurons respond, we first identified those that have spatial RFs (764 RF neurons; 21.4 +/- 1.44% of auditory responsive neurons) and plotted the RF center of each onto polar coordinates of auditory space (Figure 1D). This shows that RF neurons primarily monitor contralateral space, but some (10%) respond to stimuli from ipsilateral space. Examples of the spatial RFs of neurons that represent different areas of auditory space are shown in Figure 1C. These spatial RFs can be interpolated into the Kent distribution fit of responses to each of all 85 stimulus positions to produce a high-resolution representation of the spatial RF, which improves the accuracy of the tested models (Fig. 1C, middle). The Half Maximum Full Width (HMFW) of the Kent distribution binarizes the RF, showing the extent of auditory space it surveys (Fig. 1C, right).

**Figure 1:**
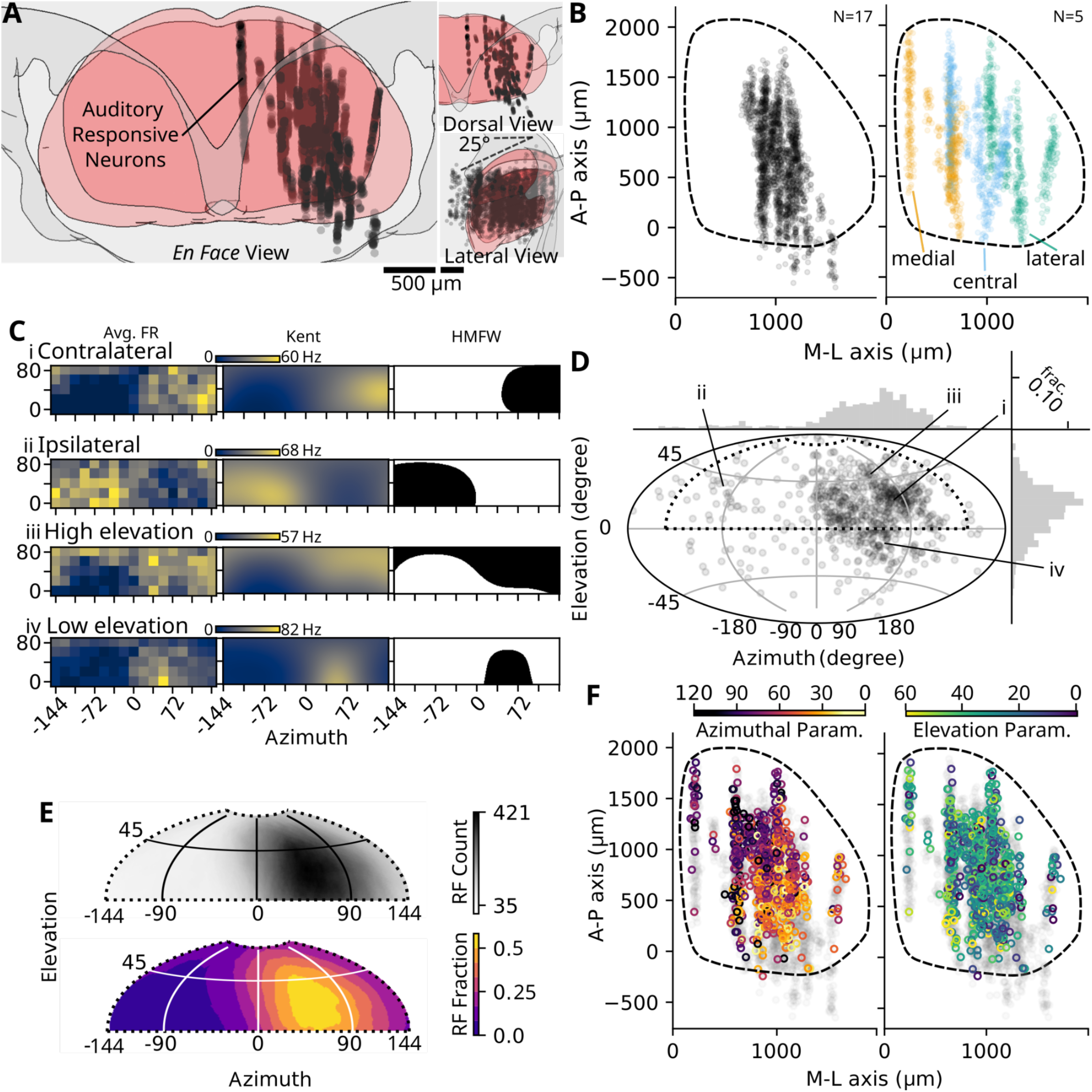
Location of the RF neurons in the SC used for modeling and the sound space to which they respond. A) Location of all auditory responsive neurons (N=32 experiments, n=3895 auditory responsive neurons) in the SC, viewed *en face* to the surface of the SC (left), or as dorsal and lateral views to show the dorsal-ventral axis of the SC. Each dot is a physiologically defined neuron, arranged along a silicon probe shank. Two scale bars at the bottom of the panels correspond to 500 μm of the two views. B) Position of all auditory neurons from single recording experiments (left; grey dots; N=17 mice, n=2054 auditory responsive neurons) and triple recording experiments (N=5 mice, n=1841 auditory responsive neurons). Each of the triple recording experiments were done within a span of 800um (400um between each insertion), within each animal there was one medial (orange), central (blue), and lateral (green) recording (total 15 experiments). C) Examples of the spatial RF of neurons that respond to sound from contralateral, ipsilateral, high elevation and low elevation space. The left graph represents the mean firing rate for each location bin, the middle graph represents the kent distribution fit the data. The two representations are on the same scale, indicated by the firing rate colorbar on the top left of the model subpanel. The right panel shows the HMFW of the kent distribution giving a binarized description of the RF, where black is the extent of the RF. D) Scatter plot of parameter fits for each RF neuron’s azimuthal and elevation position based on the kent distribution that are used in the population encoding model (n=764 neurons). Population histograms show prevalence of azimuthal parameters (top) or elevation parameters (right). Roman numerals indicate the parameter fits for examples in Ci-Civ. Black dashed region indicated is the tested area, across evenly distributed 17 azimuths and 5 elevations. Positive azimuthal values correspond to contralateral auditory space. E) Summed HMFW across all RF neurons (top, grayscale colormap). The minimum and maximum number of overlapping RF of the 764 neurons is indicated at the top of the colorbar (35-421). Contour map showing the relative fractions based on the max number of overlaps (bottom, plasma colorbar). F) Mapped contralateral azimuthal (left) and elevation (right) parameters across the surface of the SC, revealing the presence of the A-P azimuthal topographic map and an absence of the M-L elevation topographic map.

To determine the parts of auditory space that were surveilled by RF neurons, we summed the HMFW binary masks to calculate the fraction of RF neurons attending to various locations across auditory space (Fig. 1E, top), where the range of possible numbers of neurons are indicated in the grayscale colorbar (35-421). From this map, we can calculate the probability that a RF neuron has a portion of its HMFW in auditory space, as is shown in the contour map (Fig. 1E, bottom). Finally, we mapped the azimuthal and elevation parameters across the surface of the SC (Fig. 1F). In the 2D plane, azimuthal parameters created a topographic A-P map (slope: 58.98, intercept: 7.59), while elevation parameters failed to produce a M-L map, consistent with previously described analysis using all trials from the datasets (Ito et al., 2020). In conclusion, the population of RF neurons used for analysis below faithfully represents all areas of the tested auditory space, with a preference toward contralateral space.

### Statistical considerations in population encoding models

Do all neurons in our dataset contribute equally to sound localization? One method to determine how a neuron contributes to sound localization is to determine the strength of association between the auditory stimulus and the neural response by calculating Shannon’s mutual information statistic (MI, see methods). All RF neurons produce a non-zero MI, with the range from 0.02 (low amounts of information) to 0.53 (highly informative). Examples of the responses from these RF neurons are depicted as a two-dimensional heatmap and a single elevation azimuthal plot through the highest response (peak) elevation, as shown in Fig 2A i-vi; 2Ai shows a neuron with the highest MI statistic (0.53) while 2Avi shows a neuron with a low MI (0.03); four others with an intermediate MI are shown in 2Aii-v. The average MI for the population of RF neurons is 0.09 +/- 0.07 (Fig 2B, mean +/- std). To test if the MI statistic was related to anatomical mapping, azimuth, or elevation parameters, we assessed the MI distribution with respect to each of these variables (SuppFig 1). The only significant source of variation we found is that neurons associated with lower elevation fits had a larger MI, but we found no bias based on azimuthal parameter or anatomical location. We did find that the RF neurons that had a large number of action potentials and a better Kent distribution fit (i.e, smaller parameter fit errors) were associated with higher MI (SuppFig 2). Thus, MI is a single statistic that describes the relative amount of information based on the experimental task from our population of neurons; those with high MI statistics are highly responsive and well described by the Kent distribution.

**Figure 2:**
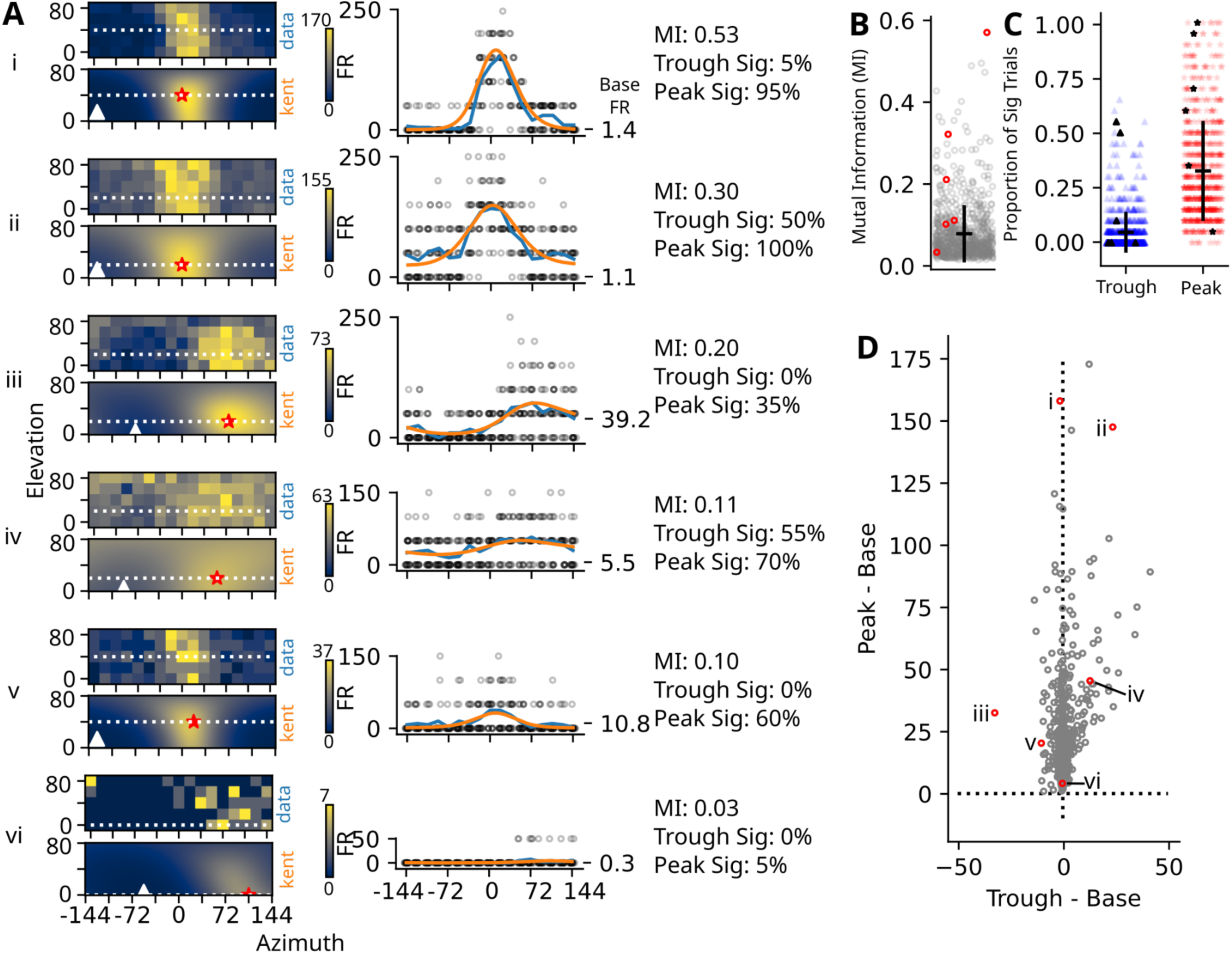
Mutual information quantifies the quality and reproducibility of trial-to-trial variation of each RF neuron. A) Six RF neuron examples (i-vi) sorted by mutual information (MI) showing trial-to-trial response variation. In each example, the two heatmaps show mean firing rate (top, data) and the model fit of the data (bottom, kent). Peak of the model fit is indicated as a red star and the trough of the model fit is indicated as a white triangle. The white dashed line indicates the elevation that passes through the peak of the RF, of which the mean dataset (blue line) and model representation (orange line) is shown in the scatter plot to the right. Each point represents an individual trial at each azimuth along the selected elevation. Baseline firing rate is indicated on the right. Far right are RF neuron descriptors of proportion of significant events based on peak or trough responses and mutual information. B) Mutual information for each RF neuron. Average and standard division are shown in black. Red data points are the 6 example neurons in A. C) Proportion of significant events based on poissonian statistics (n=764 neurons, 20 of 30 trials for model training set), using the baseline firing rate as the average (20 training trials, α=0.05 indicating significance). Proportions of significant trials for trough (blue triangles) and peak activations (red stars). Black bar represents mean and standard deviation, black data points are the 6 example neurons in A. D) Enhancement and suppression properties of all RF neurons. Scatter plot showing the trough and peak with base firing rates subtracted. Negative values suggest suppression below baseline firing rate, where positive values correspond to enhanced activity from baseline. Red data points show example neurons in A.

To further quantify the RF neuron’s responsiveness and functional behavior, we assessed the proportion of Poissonian significant trials in the testing dataset for both the highest (peak) and lowest (trough) neuronal response of auditory space, then evaluated the firing rates due to simulations from these spatial locations relative to spontaneous activity. We calculated both the proportion of significant trials in the closest experimental position to both the peak and closest to the trough, based on the Kent distribution fit (Fig 2C). We found that at the peak locations, 34.93 +/- 22.93 % of the trials were significantly different from the spontaneous rate, whereas only 4.54 +/- 10.2 % of the trough trials were significantly different from the spontaneous rate. Response patterns based on peak/trough firing rates were variable, but produced three types of patterns (Fig 2D). Some neurons have a baseline firing rate near the trough and enhanced firing rates at the peak (examples i, vi). Others have both the trough and peak firing rates elevated in response to sound, but still had spatially variant locations of activations (examples ii, iv). The third class had troughs that were suppressed below baseline firing rate but enhanced at the peak (examples iii, v). If we assume that deviation from the baseline firing rate in RF neurons indicates transmitted information, then only using the presence of action potentials (APs) should be used to inform the model. However, based on the observed characteristics of the SC RF neurons, a population decoding model that can accommodate these three types of firing patterns and the varying amounts of MI of each RF neuron may be more predictive.

### Statistical models for single-trial sound localization

To build a population pattern decoder, we used the neural responses from a single trial of auditory stimulation to estimate the stimulus location and the associated uncertainties using a weighted Poissonian likelihood model with Kent-restricted spatial RF neurons as the fundamental units.

Concatenating all the RF neurons across all the datasets produces a representation of a variety of types of RF neurons in the SC, sorted by the center of the azimuthal RF position and normalization based on RF trough to peak activity (Figure 3A). Proposed MLE and vector models used for population decoding analysis sum the weighted probability density function (PDF), or preferred direction parameter, by the firing rate (Georgopoulos et al., 1986; Jazayeri and Movshon, 2006). To replicate this in a 2D polar coordinate system, we built the model that sums the FR weighted PDF of the Kent distribution (Fig. 3B, left). This model assumes that the presence of APs, but not their absence, contains all the available location information. This model benefits by weighting neurons based on the number of APs (more information from neurons that fire a lot), and by ignoring neurons that do not produce APs, thereby eliminating the uncertainty based on the probability of firing. We assessed the accuracy of the FR model by subtracting the estimated location of the stimulus from the actual stimulus position and analyzed the accuracy along the azimuthal and elevation axes separately. Because the distribution of all the results was non-symmetric on the azimuthal prediction, we assessed the accuracy of the model for contralateral space separately from ipsilateral space. Figures 3C-D show that the contralateral but not the ipsilateral azimuth predictions using the FR model are much better than chance (mean +/- SEM, contra: 31.9° +/- 1.43°, Kolmogorov–Smirnov [KS] test p<0.0001; ipsi: 100.8° +/- 2.60°, KS test p<0.0001). Further, along the elevation, both ipsilateral and contralateral predictions were significantly different than chance (contra: 28.2° +/- 0.90°, KS test p<0.0001; ipsi: 29.7° +/- 1.00°, KS test p<0.0001), with the majority of the FR elevation predictions below chance levels in the CDF. In conclusion, the FR model works well to predict contralateral azimuthal position, but it fails to predict ipsilateral azimuthal and elevation positions regardless of location.

**Figure 3:**
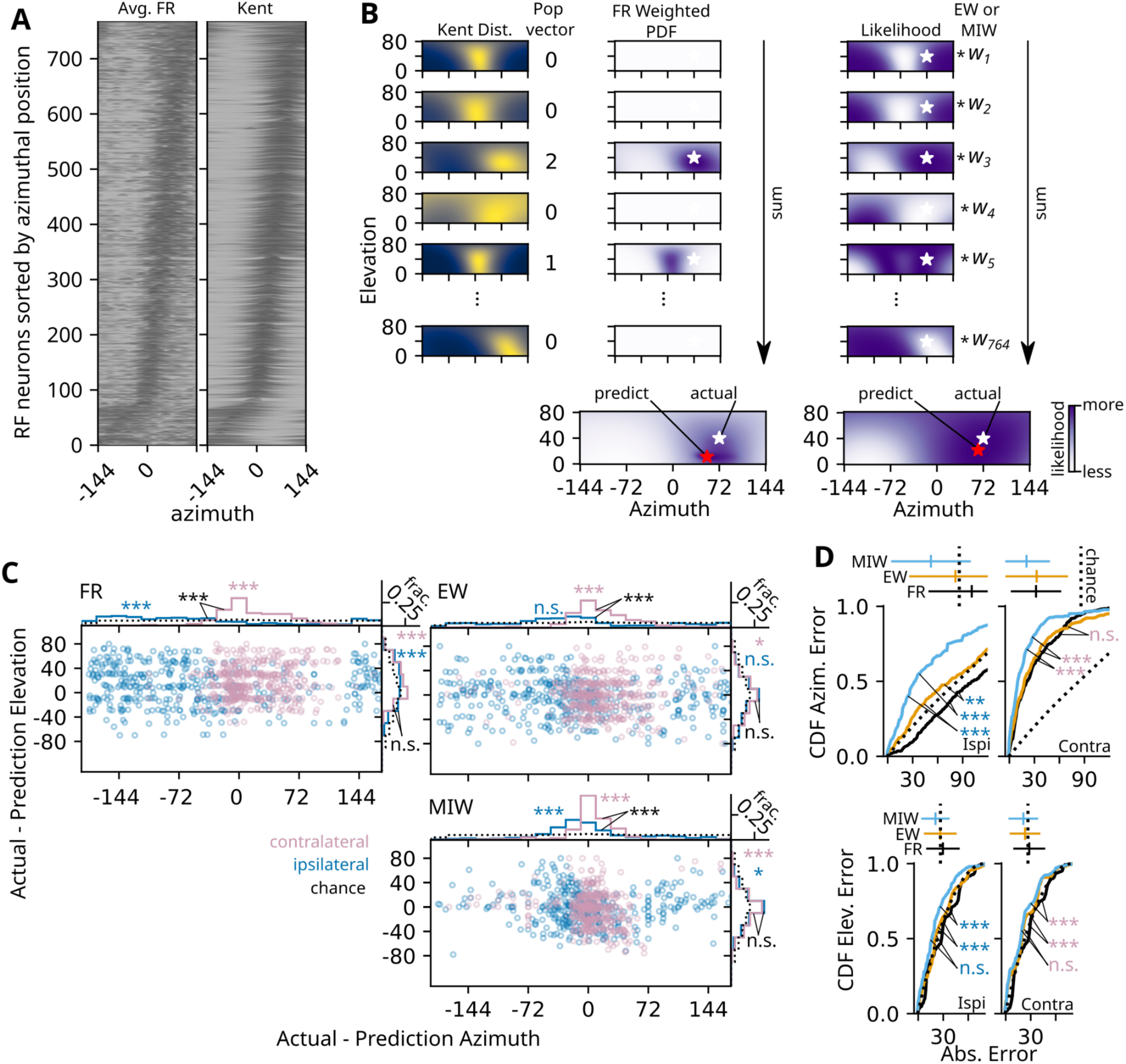
Three approaches to the summation of the weighted log-likelihoods of the population vector to predict the location of sound stimulation. A) Ridge plot of the 20° elevation of all 764 neurons selected for population encoding model sorted based on azimuthal fit. Normalized average firing rate of 20 trials (left) and the corresponding kent distribution (right). B) Schematic of the FR model (left), where the PDF of the Kent fit is weighted based on the firing rate. An example population vector, the number of AP from the first 20ms post stimulation of all neurons from a single trial, is shown to the right of the Kent fit. Schematic of the MIW and EW models (right), where the likelihood is defined by where in auditory space each number of AP in the population vector most likely to occur. The summed likelihood is weighted based on equal (EW) or mutual information (MIW). C) Results of 850 individual trials (85 positions, 10 test trials each) of azimuth and elevations for contralateral (0° to 144° azimuth, pink) and ipsilateral (-1° to - 144° azimuth, blue) auditory locations. The dashed black line corresponds to an equal likelihood of all potential results. (KS test, Bonferroni multitest correction, * p<0.05, ** p<0.001, *** p<0.0001) MIW (blue), EW (orange), and FR (black) comparison of the absolute error rates compared to chance dashed line). Cumulative distribution function (CDF) of ipsilateral (left) or contralateral (right) and azimuthal top) and elevation (bottom) error rates. Mean and standard deviation of error rates for each model shown ve CDF plots, mean chance error rate is dashed vertical line (KS test, Bonferroni multitest correction as in

To improve upon the framework of this model, we used the MI statistic to inform the model and calculated the MLE based on the number of APs from each neuron, including zero APs. The MLE model assumes that the lack of action potentials provides as much information about sound location as the presence of action potentials. To determine the sound location, we calculate the single-trial Poisson likelihood for the number of action potentials based on the Kent distribution fit (Fig. 3B right). This likelihood represents the area of auditory space from which a stimulus was most likely presented, based on the neuronal response during that single trial. In contrast to the FR model, the MLE model suggests that a majority of neurons have a higher probability to predict a stimulus location that was presented outside its own RF location. We decided to weight each likelihood based on the confidence we had in the ability of each individual neuron to predict the location; as such, we chose the MI statistics to help inform the MLE. Finally, we wanted to test if weighting based on the MI statistic benefits the model, so we built a separate model with equal weighting irrespective of the RF neurons’ response. As a result, we have two models that sum the weighted likelihood of all RF neurons based on equal (EW) or MI weights (MIW) to predict the location of the sound.

To test how weighting influences the performance of our models, we compared the accuracy of the summed log-likelihood based on the population vector using either the EW or MIW models (Fig 3C-D). Significant differences in accuracy from random distributions were found in both contralateral azimuthal predictions of EW (contra: 32.8° +/- 1.77°, KS test p<0.0001) and MIW models (contra: 20.5° +/- 0.85°, KS test p<0.0001); however, only the MIW model performed better than chance for ipsilateral azimuthal predictions (MIW ipsi: 51.9° +/- 2.39°, KS test p<0.0001; EW ipsi: 81.2° +/- 2.77°, KS test p=0.129). When we compare the azimuthal errors, we found that the MIW model produced significantly improved accuracy compared to the EW model (contra KS test, p<0.0001; ipsi KS test, p<0.0001). Further, unlike the EW model (contra: 23.2° +/- 0.86°, KS test p=0.049; ipsi: 26.5° +/- 0.98°, KS test p=0.17), only the MIW model produced significant smaller elevation error distributions from chance for both ipsilateral and contralateral positions (contra: 21.2° +/- 0.84°, KS test p<0.0001; ipsi: 20.7 +/- 0.85, KS test p=0.0072). In conclusion, model accuracy improves in azimuthal and elevation predictions when neurons are weighted based on MI that includes likelihood information from all RF neurons, not just the ones that fire in a given trial.

### The MIW model decodes azimuthal position better than the FR and EW models

In order to see how MI weighting improves the model’s performance over the FR and EW models in azimuthal prediction, we assessed how well each of the three models could predict the horizontal positions (Fig 4 and SuppFig 3). A three-way ANOVA on azimuthal error found significant sources of variation based on azimuthal position, elevation position, and model type (SuppTable1; azimuth, p<0.0001; elevation, p=0.00012 ). The overall error for the MIW model (mean +/- SEM: 32.53° +/- 1.38°, p<0.0001) outperforms the EW model (53.20° +/- 1.81°, p=0.0021); both of these performed better than the FR model (63.19° +/- 1.87°). One significant source of variance was due to interactions of elevation with model type (elevation:model, EW model p=0.0856, MIW model p<0.0001); however, interactions between models and azimuthal positions (azimuth:model, EW model p=0.112, MIW model p=0.155) and the triple interaction were not found to be significant (azimuth:elevation:model, EW model p=0.978, MIW model p=0.848). In conclusion, the MIW model significantly lowered the azimuthal error when compared to either the FR model or the EW model.

**Figure 4:**
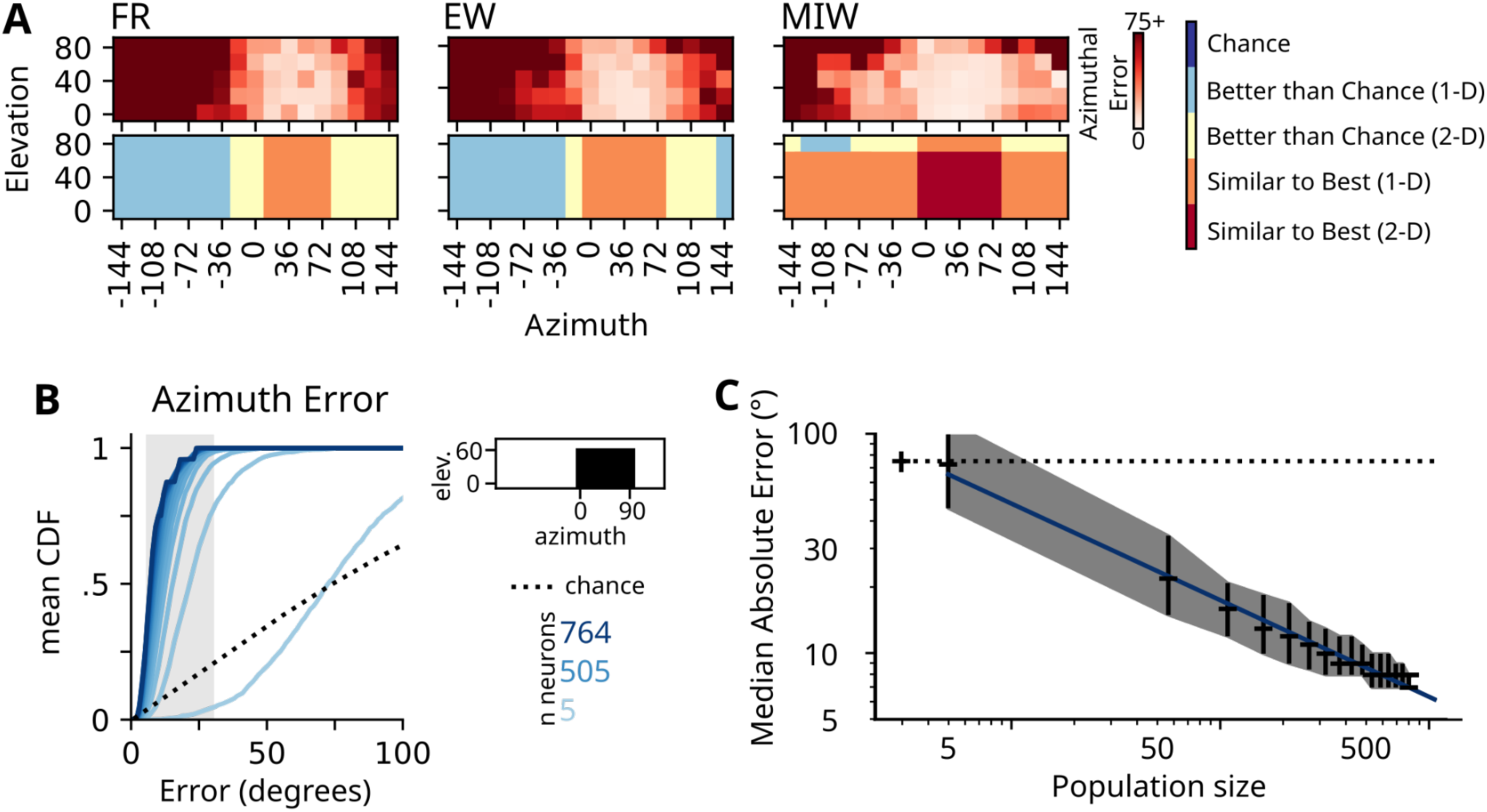
Bootstrap assessment to identify the median number of neurons necessary to identify the azimuthal location based on a single trial. A) Heatmaps showing average azimuthal error rates (red colormap, top) for FR, EW, and MIW models with indications of groups of auditory space (bottom) based on the significance shown in SuppFigure 5. B) Mean CDF of MI weighted population of azimuthal errors based on the contralateral auditory field (0° to 90° azimuth; 0° to 60° elevation; relevant area black indicated by binarized map right) with differing numbers of neurons used in the bootstrap (5 to 764; 50 neuron intervals). The light grey bar represents the range of reported behavior-relevant errors. The dashed line shows the performance based on chance. C) Log-log plot of the median absolute error of azimuthal position with respect to the population size (based on bootrapping). Bootstrapped mean and 95% CI of the median error calculated from the 100 iterations. Linear fit in log-log space (blue) to interpolate between each bootstrap.

In order to identify the elevation and azimuthal areas in auditory space that best predict azimuthal position by our models, we used collapsed errors based on azimuth and elevation to identify which positions did better than randomness, and of those, which performed the best. We assessed the better than chance performance of each model based on azimuthal position (collapsing all elevations into a single azimuthal error, Fig Supp3B) and found that all models performed well on stimuli from the contralateral side, but the MIW model was best at predicting stimuli coming from ipsilateral space. The lowest error was found at 36° azimuthal position in the MIW model with 8.06° +/- 1.14° absolute error (SuppFig 3B-C). When all errors were compared to the best performing azimuth error, we found that the EW and MIW models had better coverage of azimuthal space. We calculated the projected azimuthal errors based on elevation (collapsing all azimuths into a single elevation error rate), and found that all models and elevations performed better than chance (SuppFig 3D). The best elevation prediction was for stimuli presented at 40° elevation using the MIW model (20.54° +/- 1.88°) and similar elevation accuracy was found across 0°-60° elevations (SuppFig 3E-F). In conclusion, when mapping the lowest error rates for elevation and azimuth, the MIW model performs best in the lower (0°-60°) elevations at 0°-90° azimuths (Fig 4A, bottom).

Next, we used a bootstrapping approach to determine the minimum number of neurons needed to compute sound location based on a single trial (Fig. 4B-C). Behavioral training paradigms and psychobiological metrics were employed to determine the minimal audible angle (MAA) discrimination in the mouse. Using the auditory startle reflex as a readout, the azimuthal MAA for adult CBA/CaJ mice was determined to be between 7.5°-15° (Allen and Ison, 2010). Behavioral experiments in mice using training and water deprivation calculated an azimuthal MAA of 31° for CBA/CaJ strains (Lauer et al., 2011). Given the range in azimuthal MAA predictions, we sought to determine the number of neurons needed to achieve 6°, 18°, and 30° errors. We focused on auditory space that performed well with 764 neurons (Fig 5A-B; 0° to 90° azimuth, 0° to 60° elevation). We designed the bootstrap experiment to randomly select different RF neurons 100 times to test MIW performance across a range of neuronal counts, from 5 to 764, in 50 neuron intervals (see methods). For each of the bootstrapped counts, we show the mean cumulative distribution plot for the 100 randomly selected neurons for azimuth (Fig. 4B). As expected, we find that bootstrapping the MIW model improved azimuthal predictions with increased numbers of RF neurons.

**Figure 5.**
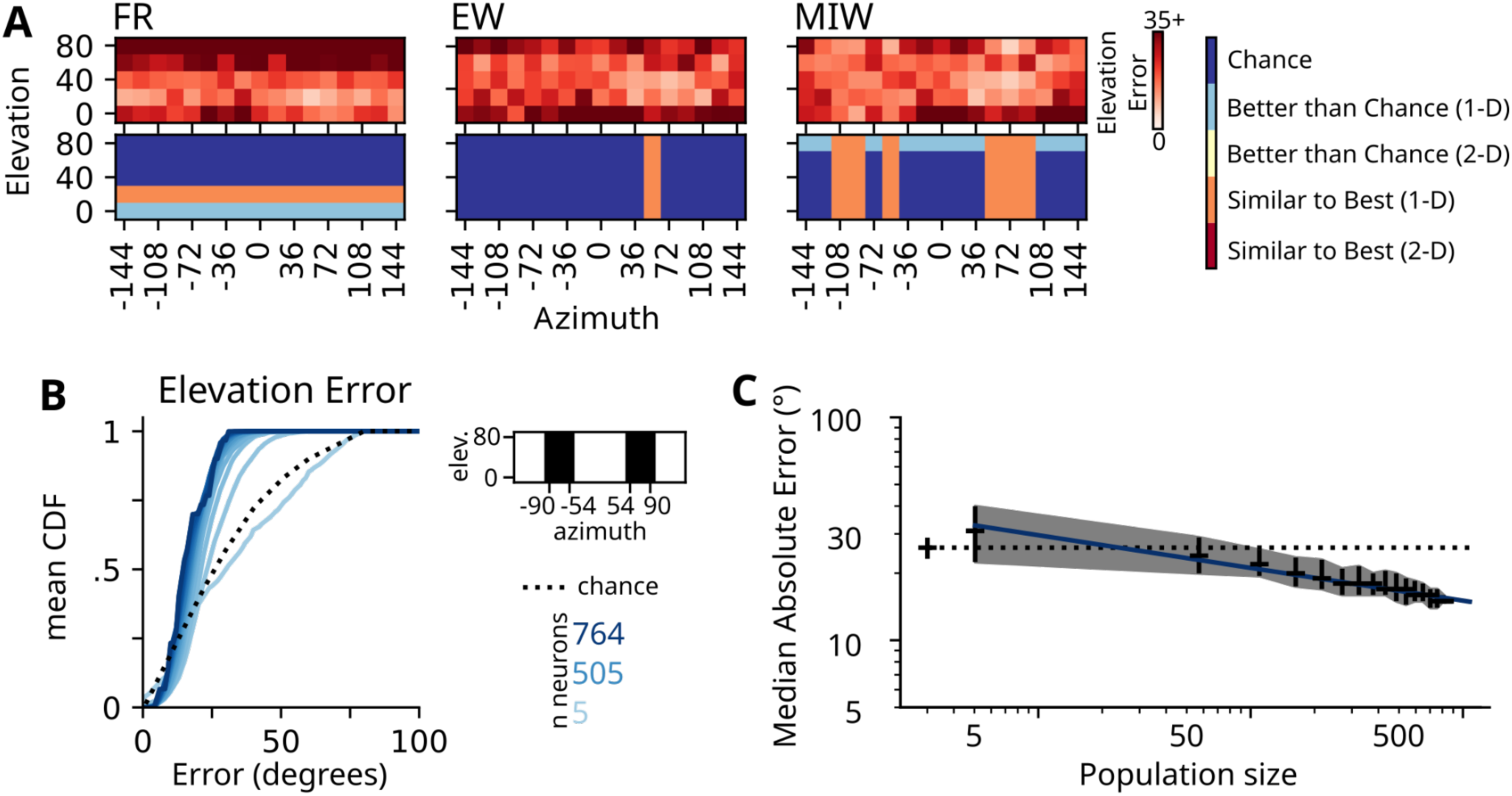
The MIW model outperforms the FR and EW models in predicting the elevation location of a stimulus in a single trial. A) Heatmaps showing average elevation error rates (top) for the three models with indications of groups of auditory space (bottom) based on the significance shown in SuppFig 6. B) Mean CDF of MI weighted population of elevation errors based on the lateral auditory field (-90° to -54°, 54°to 90° azimuth; 0° to 80° elevations; relevant area black indicated by binarized map right) with differing numbers of neurons used in the bootstrap. C) Log-log plot of the median absolute error of elevation position with respect to the population size. Bootstrapped mean and 95% CI of the median error calculated with linear fit in log-log space (blue) similar to Fig 4.

To identify the number of neurons required to achieve behavioral relevance, we quantified the minimal median error of each bootstrapped number of neurons required to achieve a given threshold. We interpolate the number of neurons necessary to achieve a given MAA by producing a linear fit (slope: -0.45, intercept: 2.13) of the log-log plot comparing the population size and the median absolute error for both azimuthal errors (Fig 4C). These graphs show that an average of 30 neurons (95% Confidence Interval (CI): 10.0-65.1 neurons) are required to achieve a 30° error along the azimuth, 92 neurons (95% CI: 42.0-171.2 neurons) for an 18° error and 649 neurons (95% CI: 477.5-920.9 neurons) to be within 7.5° of the location. In conclusion, as few as 27 auditory RF neurons are sufficient to achieve behaviorally relevant azimuthal computation of sound localization when presented in a single trial.

#### The MIW model can decode location along the elevation

We assessed how each model performs in elevation prediction by implementing a three-way ANOVA on elevation error (Fig 5 and SuppFig 4). We found significant sources of variation in elevation accuracy based on elevation position, azimuth position, and model type (elevation, p<0.0001; azimuth, p=0.0080, SuppTable2). Overall, the MIW model (20.12 +/- 0.61°; p<0.0001) performed slightly better than the EW model 24.23° +/- 0.66°, p<0.0001); both of these did better than the FR model (28.61° +/-0.68°). When we further examined the elevation error projected across elevation and azimuth, we saw that the model type was the main source of variance (elevation:model EW model p<0.0001, MIW model p<0.0001; azimuth:model EW model p=0.13, MIW model p=0.00023). Unlike the azimuthal errors, the intersection between the three variables did produce significant effects on elevation errors (azimuth:elevation:model EW model p=0.027, MIW model p<0.0001). In conclusion, elevation errors predicted by EW and MIW models are more reliable than those produced by the FR model (SuppFig 4B).

To determine the auditory space locations that best predict elevations, we used collapsed elevation errors based on azimuth and elevation to identify the groupings that did better than chance, and of those, which performed similar to the best error rate. We assessed the azimuthal positions at which the model performed significantly better than chance for elevation prediction and found that lateral azimuthal positions in contralateral (58°-90°) and ipsilateral (58°, 90°-108°) auditory space were significant (SuppFig 4F). These results align with the first order linear fits on the actual vs predicted plots (SuppFig 4E). All azimuth positions that predicted better than chance had errors similar to the best error, located at the contralateral 72° azimuthal position (14.20° +/- 2.26°). We then determined that the lower elevations (0°-20°) in the FR model and the highest elevation (80°) in the MIW model were predicted better than chance. The lowest error was from the FR model, at 20° elevation position (14.20° +/- 0.76°); however, this is because this model has mostly predictions at lower elevations (SuppFig 6A). Thus, we found that the MIW model was the only model to predict elevation at lateral azimuthal positions across the five tested elevations better than chance.

**Figure 6:**
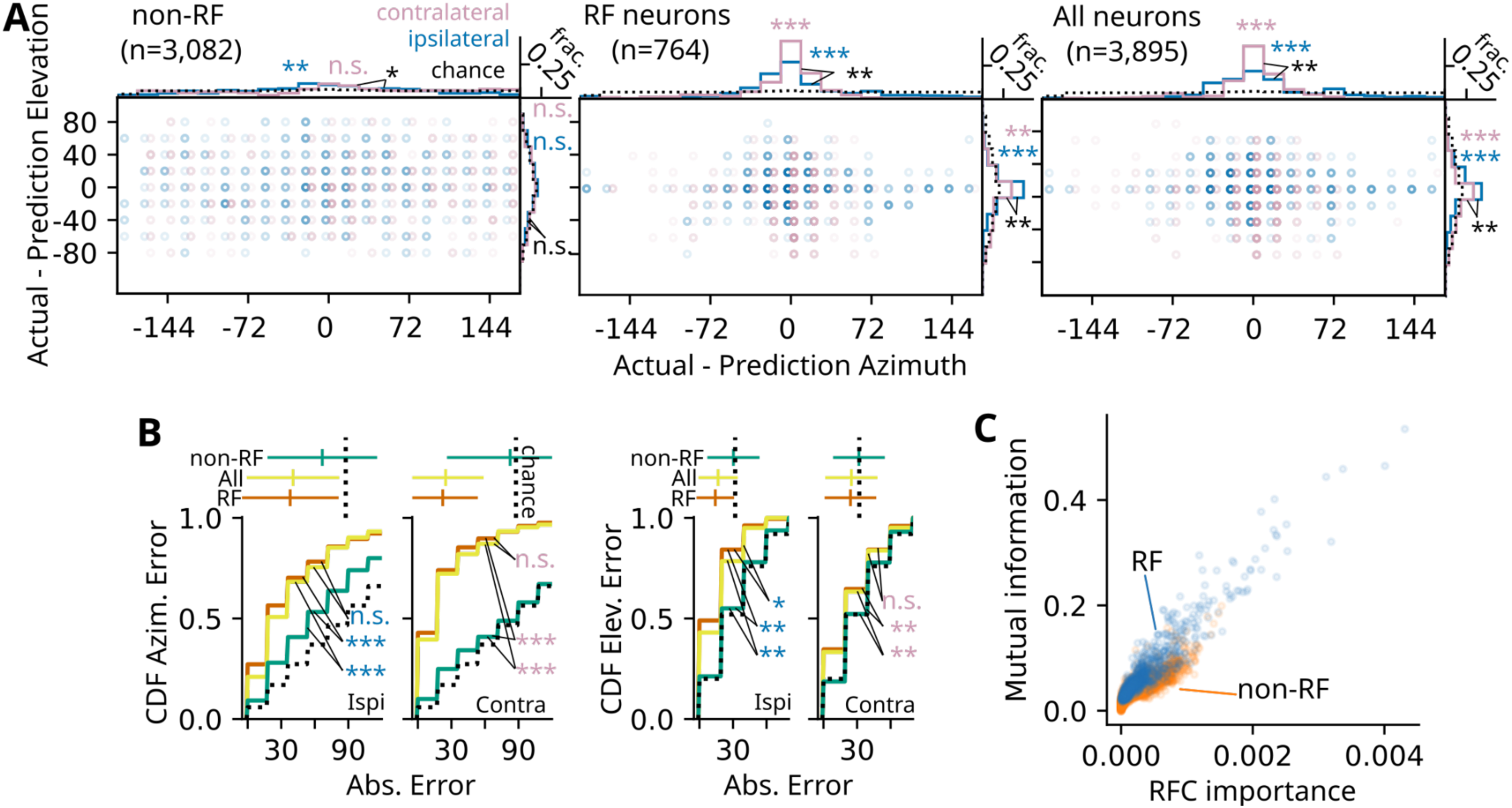
Sound localization information in the SC comes from RF neurons. A) Error plot of actual minus prediction indicating elevation and azimuthal accuracy for contralateral (pink) and ipsilateral (blue) auditory locations from Random Forest Classifier (RFC) models trained with Non-RF, RF neurons, or all neurons. B) Absolute error rates of RFC trained on RF (orange), All (yellow), and non-RF (blue-green) neuronal grouping compared to chance (dashed line). CDF of ipsilateral (left) or contralateral (right) and azimuthal (far left) and elevation (far right) error rates. Mean and standard deviation of error rates for each model shown above CDF plots, mean random error rate is dashed vertical line (KS test, Bonferroni multitest correction as in A). C) Scatter plot from all neuron RFC showing the relationship between the importance metric, based on the Gini Impurity, and the calculated Mutual Information index.

To identify the number of neurons required to achieve behavioral relevance, we quantified the minimal median error of each bootstrapped number of neurons to achieve an 18° error (a 30° error is achievable based on chance). Lateral auditory space (-90° to -58°, 58° to 90° azimuth, 0° to 80° elevation) was the only location able to achieve elevation predictions (Fig 5A-B). Using a similar bootstrapping technique as in figure 4B-C, we interpolated the number of neurons necessary to achieve a given MAA by producing a linear fit (slope: -0.15, intercept: 1.62) of the log-log plot comparing the population size and the median absolute error for both elevation errors (Fig 5B-C). We find that 283 neurons (95% CI: 94-598.6) are required to predict elevation localization within 18°. In conclusion, to achieve the same error rate, elevation prediction from the MIW model requires substantially more neurons than azimuthal prediction.

#### RF neurons contain the auditory location information in the SC

To test the assumption that RF neurons are the only neurons that contain spatial information in the SC, we trained a Random Forest Classifier (RFC) on all auditory responsive neurons, RF neurons, and non-RF neurons (Fig. 6). A RFC is an ensemble machine learning tool that utilizes decision trees for multi-label applications (Chaudhary et al., 2016) where, in this case, elevation and azimuthal predictions are established on the same training/testing session. Similar to the MIW model, the algorithm uses either the Gini Impurity index or Entropy as the criterion to simplify decision trees for the ensemble. Using entropy as the criterion ultimately weighs the classification decisions more heavily on informative features, improving generalizability (Quinlan, 1996, 1986). However, unlike the likelihood model, the classifier does not require spatial restriction modeling in the framework; thus, any neuron can be utilized to train and test the model. Therefore, we determined the cumulative distribution of the predicted subtracted from the actual positions along both azimuth and elevation using only non-RF neurons, RF neurons, or all neurons. We found that the RF neuron based model predicts sound location better than that based on non-RF neurons (Fig. 6A-B). Specifically, the predictions of azimuthal location using RF neurons were significantly better than chance for stimuli presented from ipsilateral or contralateral auditory space (RF ipsi: 38.0° +/- 2.17°, KS test p<0.0001; RF contra: 22.2° +/- 1.48°, KS test p<0.0001) and along the elevation (RF ipsi: 23.1° +/- 1.09°, KS test p=0.000124; RF contra: 14.2° +/- 0.84°, KS test p=0.000702). However, when using the non-RF neurons in the model, the contralateral azimuthal prediction only significantly better than chance for ipsilateral stimuli (non-RF ipsi: 66.7° +/- 2.45°, KS test p<0.0001; non-RF contra: 82.5° +/- 2.66°, KS test p=0.967). The elevation predictions were also not above chance (non-RF ipsi: 30.4° +/- 1.17°, KS test p=0.974; non-RF contra: 31.6° +/- 1.09°, KS test p=0.999). Furthermore, all locations were better predicted in the model that used RF neurons than one that used non-RF neurons (non-RF v RF ipsi azimuth KS test, p<0.0001; non-RF v RF contra azimuth KS test, p<0.0001; non-RF v RF ipsi elevation KS test, p=0.000194; non-RF v RF contra elevation KS test, p=0.00100). These results are consistent with the hypothesis that RF neurons contain the majority of spatial information in the SC.

To discover if including non-RF neurons in the model improves the prediction accuracy from the RF neurons alone, we trained the model with all neurons and compared the predictions with those from the RF neuron model. We found that there were minor improvements in the accuracy of the model (Fig 6A-B) (All v RF ipsi azimuth KS test, p=0.416; All v RF contra azimuth KS test, p=0.939; All v RF ipsi elevation KS test, p=0.0366; All v RF contra elevation KS test, p=0.939). The RFC allows us to examine which neurons were most informative to the model based on the Gini Impurity. As expected, the Mutual Information statistics were similar to the RFC importance, since entropy was used as the criterion in training the model (Fig. 6C). In conclusion, we show that including all auditory neurons failed to improve the predictions of sound localization from a model trained on only RF neurons.

## Discussion

In this paper we used previously described data sets obtained from in vivo recordings of mouse SC neurons, in response to spatially localized white noise stimuli, to determine the ability of the SC to identify the elevation and azimuthal location of a sound source in a single trial. The model that works best is a Mutual Information weighted Poissonian likelihood model that is based on the number of APs for a given trial, thereby summing across the full population of identified RF neurons to determine the sound location. One major assumption this model makes is that all the spatial information is based solely on RF neurons. We test this assumption with a Random Forest Classifier and show that including all auditory responsive neurons does not improve sound source prediction. Based on the population of neurons described in this paper, we tested the sensitivity of this model and found that, on average, 30 SC RF neurons are required to determine the location of a sound to behaviorally relevant accuracy.

Two main models used to determine auditory localization abilities of a population of neurons are rate code based models and population/place code models, and each has been used to model the ability of a population of neurons to localize a sound source in different auditory areas of the brain. Rate-code strategy models use the differences in firing rates of neuronal pairs or full hemispheric comparisons (opponent channels) to calculate the location of a sound source, and have been used to model the ability of neurons in the IC and auditory cortex to encode sound location (Groh et al., 2003; Lesica et al., 2010; McAlpine et al., 2001; Miller and Recanzone, 2009; Ortiz-Rios et al., 2017; Panniello et al., 2018; Stecker et al., 2005). Population, or place code models, account for the diversity of spatial responses within a population of neurons to predict sound location, typically using spatially restricted responses (Belliveau et al., 2014; Day and Delgutte, 2013; Fischer and Peña, 2011; van der Heijden et al., 2019). Population codes are more informative of sound location than rate codes; however, functional modeling studies from a variety of species show the brain has the ability for rate coding (Belliveau et al., 2014; Fischer and Peña, 2011; van der Heijden et al., 2019). Finally, hybrid models based on neuronal responses in the IC have been proposed, acknowledging that the two computations can cooperate in sound localization (Chen and Song, 2024).

Because of the existence of auditory RF neurons in the mouse SC, we chose to use a population code model to compute sound source location when presented with a single trial. As such, we built an MLE model based on the RF neurons, summing the likelihood of sound localization of each RF neuron. We compared three models with different approaches to the final computation: Firing rate (FR) weighting of the PDF, Equal (EW) or Mutual Information weighting (MIW) of the Poissonian likelihood based on the number of APs for a given trial. Mathematically, the MI weighting gives the model more algorithmic credence to RF neurons that are more reliable in firing. All three models did relatively well when assessed at the 0°-72° azimuthal locations; however, the MIW was superior to both models in azimuth predictions outside of this range. In addition, the MIW was the only model to achieve predictions better than chance for elevation.

One major assumption of these models is that all the spatial information is based solely on RF neurons. We tested this assumption by training a Random Forest Classifier (RFC) on all auditory responsive neurons, both RF neurons and non-RF neurons (Fig 6). Previous models have found that auditory responsive neurons that do not have spatially restricted RFs can contribute to sound localization (Mazo et al., 2024; Groh et al., 2003; Lee and Groh, 2014; Werner-Reiss and Groh, 2008). Our results show that the responses from non-RF neurons were unable to predict the location of a stimulus, and that combining all auditory neurons with the RF neurons did not improve the sound localization predictions of the model trained on only RF neurons. Taken together, these results confirm that sound source location can be modeled with the population response of a set of SC RF neurons.

We find that the MIW model can predict the location of a sound within behaviorally relevant accuracy. A few studies that utilized different mouse strains and auditory behavior assays have reported MAAs between 7°-30° for azimuthal localization and 80+° in the medial plane (azimuthal position 0°) for elevation (Allen and Ison, 2010; Lauer et al., 2011). A 30° MAA has also been used as a behaviorally relevant cutoff in population models that use IC sound responses to predict azimuthal location (Boffi et al., 2024). Therefore, we asked how many RF neurons are needed to estimate locations within these ranges using the MIW encoding model. We show that, along just the contralateral azimuth, on average, 30 neurons are required to predict the location of a stimulus within a 30° azimuthal error, 92 neurons to predict within an 18° error, and more than 649 neurons to predict the location with a 7° error (Fig 4). For elevation stimulus location, the model worked best at predicting sound location at peripheral contralateral and ipsilateral locations between 58°-90° azimuth (Fig 5). We found that on average, 283 neurons were required to achieve an 18° elevation error. Even though ipsilateral sound space had far fewer representative RF neurons, the elevation prediction had similar accuracy as that obtained from contralateral space. This accuracy is also similar to what has been observed behaviorally. Although our model could not predict elevation better than chance at the 0° azimuth, the elevation MAA of 80+° was determined at this location (Lauer et al., 2011). Our results compare well with studies using population encoding models in the IC of different animals. Based on recordings from the rabbit IC, 21 neurons are required to localize a sound within a MAA of 15° (Day and Delgutte, 2013). Similarly, modeling of auditory responses in the owl optic tectum showed ∼40 neurons could achieve an azimuthal error of 10° (Fischer and Peña, 2011).

In conclusion we find that mouse SC neurons that contain spatially restricted RFs can use population coding strategies to localize sound in a single trial and the localization is better along the azimuth than the elevation. This is consistent with the few studies that have measured mouse MAAs, and correlates with the data that shows mice have a topographic map of auditory space along the azimuth but not elevation (Ito et al., 2020). The correlation between topography and spatial localization ability is consistent with the hypothesis that one feature of topography is to help with spatial/temporal processing (Kaas, 1997), although this idea has been challenged (Avitan et al., 2016; Weinberg, 1997). One limitation of our analysis is that it relied on a data set obtained from the recording of mice passively listening to (50db) white noise bursts of sound. It has been shown that auditory RF neurons can have a broadening of the RF or different elevation responses based on loudness, whereas azimuthal position is unaffected (Remington and Wang, 2018). Attention can also change the response properties of RF neurons (Hu and Dan, 2022), and subtle movements of the ears could lead to errors in our measurements. Our analyses also reveal that there are at least 3 types of responses of SC neurons to spatially restricted stimuli, differing in their relative responses to stimuli outside their RFs; some fire similar to their spontaneous rate, some suppress below their baseline levels, and others have an elevated firing rate above the baseline outside the RF, but still had spatial preference (Fig. 2). The significance of these differences remains to be understood. However, combined with previous analysis that shows 4 types of temporal responses of auditory SC neurons to white noise (Si et al., 2025) this data may reflect the need to localize sounds of varying spectral-temporal and intensity properties. Therefore, further dissecting the relationship between these neuronal types and context dependent sound localization is an important direction of future research.

## Methods

### Data

The SC neural response data used to build and test the models was obtained from 4-shank, 256 channel, silicon probe recordings from head-fixed CBA/CaJ mice, freely running on a cylindrical treadmill, that have been previously reported (Ito et al., 2021, 2020). We concatenated two previously described datasets to build and test different population decoding models to have a large number of SC neurons with spatially restricted RFs that represent all anatomical and auditory spatial positions. This resulted in 17 single penetration experiments (N=17 experiments, N=17 mice) and 5 triple penetration experiments (meaning that the probe was moved to three distinct locations in the same mouse, N=15 experiments, N=5 mice). All experiments that were concatenated were done using 30 trials of white noise (5-80 kHz) from 85 sound locations; 17 azimuths (-144° to 144°, 18°intervals) and 5 elevations across the upper auditory field (0° to 80°, 20° intervals). The data utilized in this manuscript are single-unit neurons identified through the lab’s custom-designed spike sorter, as previously described (Ito et al., 2017; Litke et al., 2004). We chose to focus on the first 20 ms after the response, as most RF neurons have a peak response within 20 ms after the stimulus onset; therefore, we restricted our population decoder by using just the responses of RF neurons in this time window. Approximately 20% of auditory responsive neurons have RFs, determined as such if their responses were better described by the Kent distribution than the uniform distribution (see below). This report only uses the spike-sorted data, without any previous descriptors. All data in this manuscript was processed using a custom Python script that will be shared on the Feldheim lab GitHub page (https://github.com/feldheimlab/RF-population-encoding).

#### Axonal vs somal waveform

We found that we had two classes of waveforms from RF neurons: biphasic and triphasic (SuppFig 5). Triphasic waveforms are thought to originate from myelinated axons or neurites (Bakkum et al., 2013; Barry, 2015; Robbins et al., 2013). We excluded these signals from the population models because we wanted to include only SC originating neural spikes.

#### Splitting and data quality checks

For model testing we determined which neurons had spatially restricted receptive fields that originated in the SC, based on their responses to 20 of the 30 trials at each of the 85 positions, and we set aside the remaining 10 trials to test the models. For all data descriptions, model fits, and weight approximations, the model only used randomly selected 20 trials at each experimentally tested location; the remaining 10 trials were used to test the model’s performance and were not used to inform the descriptions. All data presented in this manuscript and characterizations are similar to what was previously reported, when all 30 trials were included in their characterization. The breakdown of all experimental data, including the number of RF neurons and the shifts in population based on insertion location is characterized (SuppFig. 6).

#### Ridge plot

The average firing rate and the Kent distribution were normalized with the same factor, where we subtracted off the trough firing rate (making this 0) and set the peak equal to 1. The RF neurons were sorted based on the preferred location of azimuth. The 20° elevation was chosen because it showed the most diversity of all RF neurons (Fig 3A).

### Anatomical mapping of auditory responsive neurons

Metadata, descriptors of the experiment which include location and the depth of probe insertion from each recording, were used to map experiments to the corresponding location in the SC. We modified the 2-D mapping (depth and either M-L or A-P) of the recorded neurons in the SC previously described in (Ito et al., 2021; Ito et al., 2020) to include 3-D plots of auditory and RF neurons. From the metadata, the probe insertion depth, relative movement of shank position for multi-insertion experiments, and the visual RFs mapped for each of the shanks and the position of the front-most shank that was implanted into the SC was determined. Once the position of the front shank was determined, we restricted the rigidity of the probe and approximated the angle of insertion based on the visual responses of the other shanks, and rotated the probe accordingly. For the triple insertion mice, we did the same approximation for the center insertion and had metadata on the relative distances between each penetration. We assumed the angle of insertion was the same for each of the three experiments. From this metadata, the 3-D map of all auditory neurons was done with Brainrenderer for all experiments (Claudi et al., 2021).

### Receptive field neurons

#### Auditory responsive neurons

Auditory responsive neurons were identified as previously described (Ito et al., 2020). In short, the first 20ms post-stimulus response was used to calculate the probability that the response was elevated shortly after the auditory stimulus, based on quasi-Poisson statistics of the baseline firing rates. Auditory responsive neurons were identified based on the significance tests such that the neuron’s spike count had a p-value less than 0.001. Of the 6025 single unit recordings, 3895 were auditory responsive (64.6%).

#### Kent distribution fit

The probability density function for the Kent distribution is defined as the following:

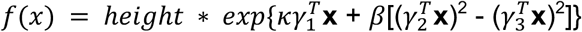

Where **x** denotes 3-D cartesian coordinates, *κ* and *β* are the parameters for concentration and ellipticity of the area, respectively. *γ* is the spherical unit that is based on the polar coordinates of the azimuth (*θ*) and elevation (*ϕ*) with a rotation parameter (*α*) of that unit. Instead of computing the normalizing constant based on the *κ* and *β* for the probability density, we fit a height parameter to ensure the probability distribution is scaled to the number of action potentials present in the data (see Fig 3). While the Kent distribution is fit in Cartesian coordinates, we report all location parameters with polar coordinates. Functions were written in python, utilizing python’s scipy optimization packages (Virtanen et al., 2020).

All auditory responsive neurons were fit with both a Kent and a uniform distribution. The Bayesian information criterion (BIC), as defined by the Residual Sum of Squares (RSS), is defined as:

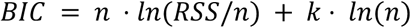

Where n is the number of observations and k is the number of parameters to fit the model. RF neurons were identified by comparing the BIC between the two distributions. Those neurons that had better BIC based on the Kent distribution than the uniform, were defined as receptive field neurons.

We interpolated the Kent distribution to allow us to get a more precise representation of the location accuracy of the model, meaning that we did not restrict the predicted value to the 85 experimentally tested positions. We interpolated within the tested area (Azimuth [-144,144], Elevation [0,80]), with 1 degree spacing in each direction. This assumes that the neuron would respond in a similar spatially restricted fashion as the Kent distribution fit between the tested locations.

#### Full width half maximum (FWHM)

The FWHM was determined by thresholding the interpolated Kent distribution at the halfway point between the minimum and maximum of the Kent distribution. Given the variety of response properties, we could not just subtract out the baseline stimulus but based it on the response to stimuli. This was necessary given that some neurons were elevated at every location, but still had a spatial preference.

The sum of all FWHM receptive fields was then divided by the number of RF neurons used to calculate the distribution across auditory space. Assuming we have a decent representation of RF neurons, we can quantify how many neurons attend to different areas of auditory space. Importantly, there was never a space of auditory space that was lacking in the representation of RF neurons.

### RF neurons statistics

#### Mutual information

The mutual information statistic (Nelken and Chechik, 2007; Panzeri et al., 2010) is defined as:

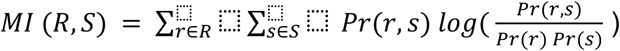

The mutual information was calculated based on the probability of stimuli and neuronal responses across the 85 positions and 20 trials.

#### Peak and trough significance

Peak (highest activity in the RF) and trough (lowest activity in the RF) were found based on the fit kent distribution. The closest actual stimulation site was used to calculate the Poissonian significance (*α* = 0.01) from spontaneous activity for the 20 trials used for the kent fit. Percent of significant trials were determined by dividing the number of trials that were significant from spontaneous by the number of total trials. Spontaneous activity for each neuron was determined by averaging all trials at 1980-2000 ms after the stimulus was presented, well after midbrain auditory processing was complete.

### Population encoding models

#### Firing rate (FR) model

The model prediction is based on the FR weighted sum of kent probability density function (PDF):

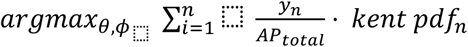

The kent PDF was calculated from the fit kent distribution above, normalized by dividing the sum of all values in the kent fit. The weighting of the firing rate was normalized to the total number of action potentials (*AP*_*total*_) in the population vector for that trial, thus the sum of all FR weights equals 1. *y* is the population vector for a single trial, n is each neuron in the population encoding model. Therefore when the number of AP for that neuron is 0, the PDF of that neuron does not influence the prediction. We sum the each PDF to achieve the maximal likelihood, selecting the most likely for both azimuthal (*θ*) and elevation (*ϕ*) positions.

#### Mutual Information and equal weights (MIW and EW)

Weights for mutual information were normalized based on the sum of mutual information statistics for all neurons included in the model, such that the sum of all weights is equal to 1. Equal weights are defined as 1/n, where n is the number of neurons included in the predictive model.

#### Summed log-likelihood classifier

The model prediction is based on the weighted sum of the Poisson log-likelihood:

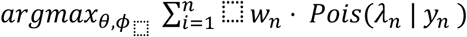

where *λ* = *f*(*x*) which is defined by the spatially restricted Kent distribution, *w* is the corresponding weight, *y* is the population vector for a single trial, and n is the specific RF neuron in the model. Given that the interpolated area of the Kent distribution is a 2-D array, we can predict both the azimuth (*θ*) and elevation (*ϕ*) location from the summed likelihood of a full population vector single trial.

#### Random Forest classifier

A Random Forest Classifier (RFC) from the Python scikit-learn package (“Machine Learning in Python, Pedregosa,” 2011) was used in supervised learning to predict the sound localization based on a single trial. A 20 / 10 trial split was used to train and test each of the three models, based on RF neurons only (n=764), all auditory responsive neurons (n=3895), or non-RF neurons (n=3082). RFC allows for multilabel classification, but we were only able to train based on the 85 tested positions and could not interpolate between the locations. Stratified splits, based on stimulus location, ensured that we had evenly distributed training and testing datasets across all 17 azimuths and 5 elevations resulting in 1700 total population vectors were used for training, and 850 total for testing for each model. Each model was trained and tested on the same subset of data.

### Statistical tests

#### Linear regression

We chose a linear fit to assess how well the model predicts, based on the unity line. In this analysis, we assess the performative prediction of the model, where we use the *R^2^* values to assess the closeness to the fit line, where the higher the value reflects the better the linear fit explained by the data.

#### Kolmogorov–Smirnov

Kolmogorov–Smirnov (KS) tests were used with Bonferroni multiple-test comparison correction to determine distinct cumulative distributions. These statistics were run from the scipy Python packages (Virtanen et al., 2020). The random distribution was determined based on the distribution of all possible outcomes of the comparison.

#### 3-way ANOVA

An analysis of variance (ANOVA) was performed by comparing the fits of an Ordinary least squares (OLS) linear model, which assumes normality, and Generalized Linear Models (GLM) using either the Gamma distribution ( with a log link function) or the Tweedie distribution (hyper parameter var power set to 1.5 and Log link function, which corresponds to a compound Poisson-Gamma mixture model) from statsmodels (Seabold and Perktold, 2010). When we compared the AIC and BIC of these three models, the Tweedie distribution was a better fit over the two others. Given the diversity of model types, elevation, and azimuth, the thee-way ANOVA model best suited for overdispersion seems to have been the tweedie distribution.

When determining the significance of the coefficients using the GLM, we used the Wald statistic (z-value) that quantifies the number of standard errors between the parameter estimate and zero, where negative z-value relates to a negative coefficient (SuppTable 1-2). We interpret significance if that coefficient has a p-value is less than 0.05, such that the predictor coefficient has significant impact on the change in errors. The categorical coefficients of model types, all values are done in comparison to the common FR model. For post hoc analysis to determine better than chance or similar to best, we utilized KS tests as they do not assume normality.

#### Bootstrapping

Determining the number of neurons necessary to achieve accuracy of sound localization was done utilizing bootstrapping. We randomly selected 5 to 755 neurons, with 50 cell intervals, resulting in 16 iterations of testing the model with varying numbers of neurons. Each round was assessed 100 times and the mean and 95% Confidence Interval of each iteration was reported. This allows us to determine the distribution of accuracies that would arise from the population of cells described in this study.

#### Interpolation of bootstrapping results

The population size (bootstrapping iterations) and the median absolute error from each iteration was used to calculate the mean and standard error of medians. They are represented in a log-log plot, where we fit a linear model in log-log space of the mean and errors of the mean (Fig 6). We are able to interpolate the number of neurons required to achieve various behaviorally relevant errors, based on this linear fit.

## Acknowledgements

This work was supported by the Brain Research Seed Funding provided by University of California, Santa Cruz, the National Institutes of Health Grant 1R01DC018580 to D.A.F, and a NIH/NICHD T32 Training Award T32HD108079 to B.R.M. We thank Jena Yamada, Greta Vargova, Gursajan Gill, and Veronica Ostrin for their helpful comments on the manuscript.

## Supplemental Figures

**Supp Figure 1:**
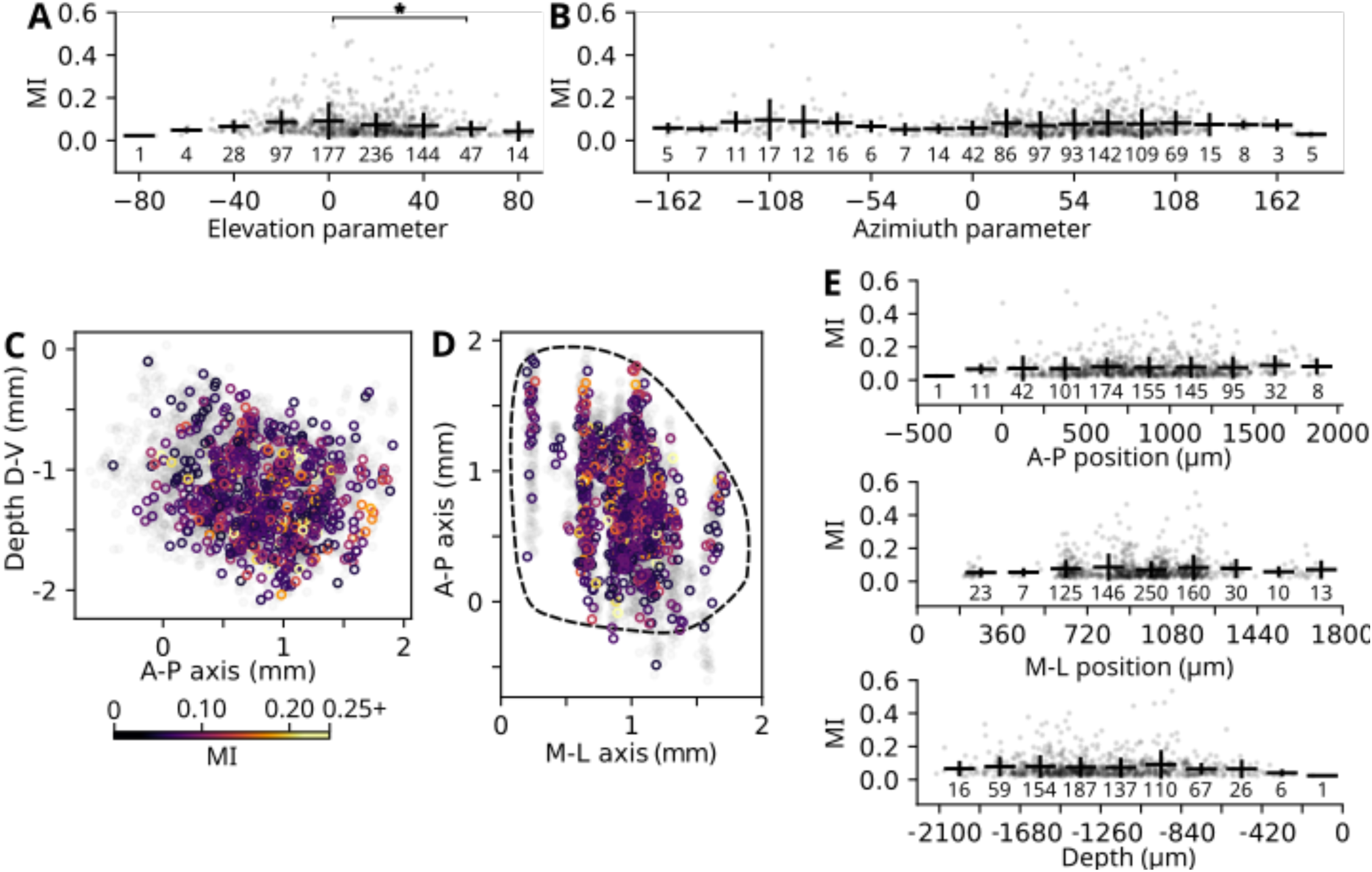
The MI statistic does not have apparent bias/preference in parameter or anatomical space. A) Mutual information (MI) plotted by elevation parameter. Each MI value was placed in a 20**°** bin across the tested space, and the average and standard deviation of the binned values are reported. (F statistic:3.149, *df*=8, *p-value*=0.0016). Tukey post-hoc analysis (* p<0.05), showing a slight bias for higher MI in neurons that have parameter fits in lower elevations, showing significant difference between 0° and 60°. B) MI plotted by azimuthal parameter, similar to A but each MI value was placed in a 18**°** bin. (F statistic: 0.7805, *df*=19, *p-value*=0.732) showing no difference in MI distribution based on azimuthal parameter fits.. C) MI of each RF neuron plotted with respect to depth and A-P axis. D) MI plotted in the *en face* anatomical view. E) MI plotted by anatomical positions, showing no significant localization of MI based on binned MI values and an ANOVA. Top plot shows A-P position with 250um bins (F statistic:0.5171, *df*=9, *p-value*=0.8628), middle plot shows M-L position with 180 um bins (F statistic:1.4634, *df*=9, *p-value*=0.1573), and the bottom plot shows depth position with 210 um bins (F statistic:1.3840, *df*=8, *p-value*=0.1997). Showing no difference in MI neurons based on anatomical position.

**Supp Figure 2:**
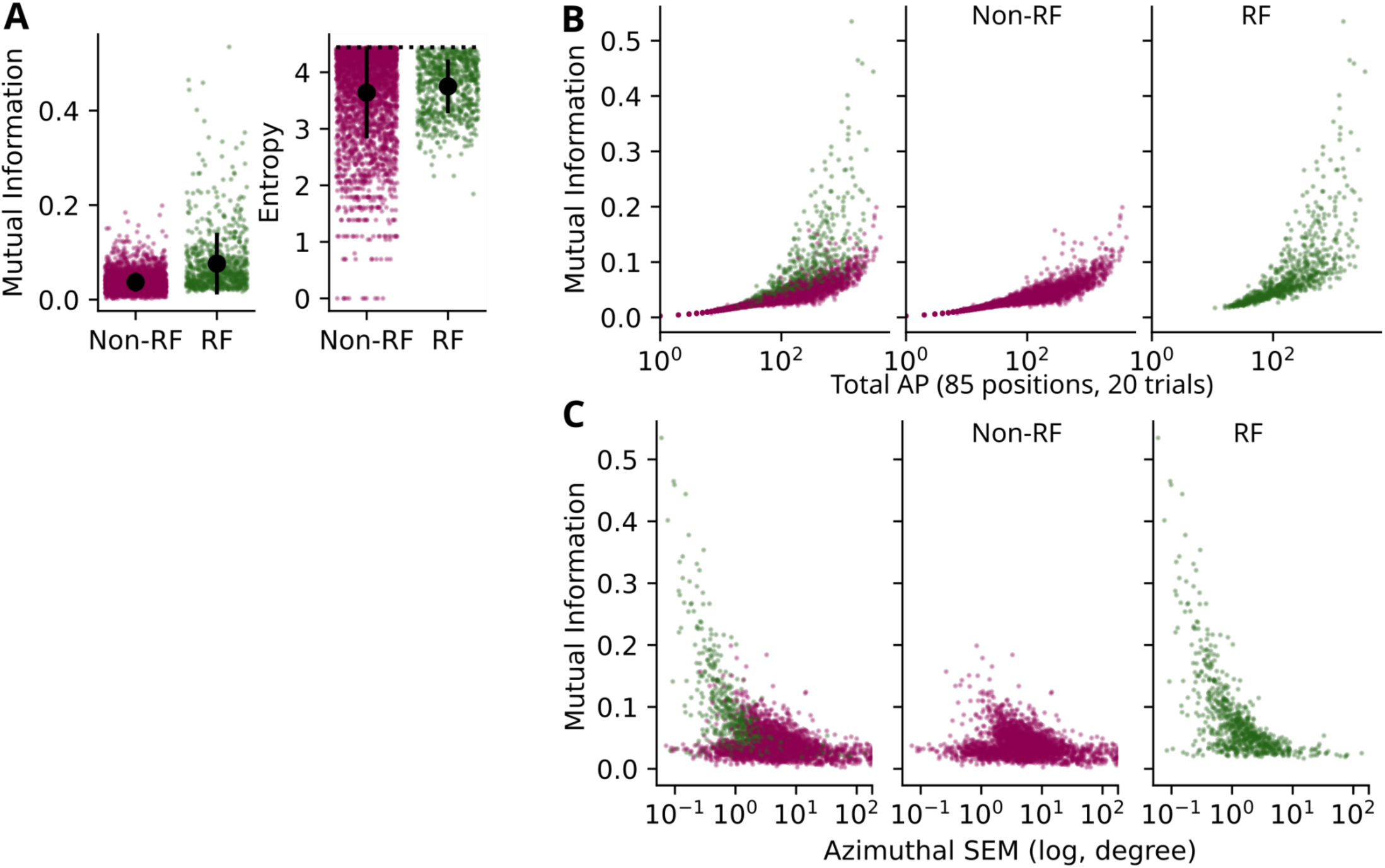
RF neurons contain more information than non-RF neurons. A) Mutual information and entropy metrics for non-RF auditory neurons (pink) compared to RF neurons (green). B) Mutual information plotted by the number of APs used to calculate the metrics for non-RF auditory neurons (pink) compared to RF neurons (green). C) Azimuthal parameter standard error of the mean (SEM) plotted by total number of APs for non-RF auditory neurons (pink) compared to RF neurons (green).

**Supp Figure 3:**
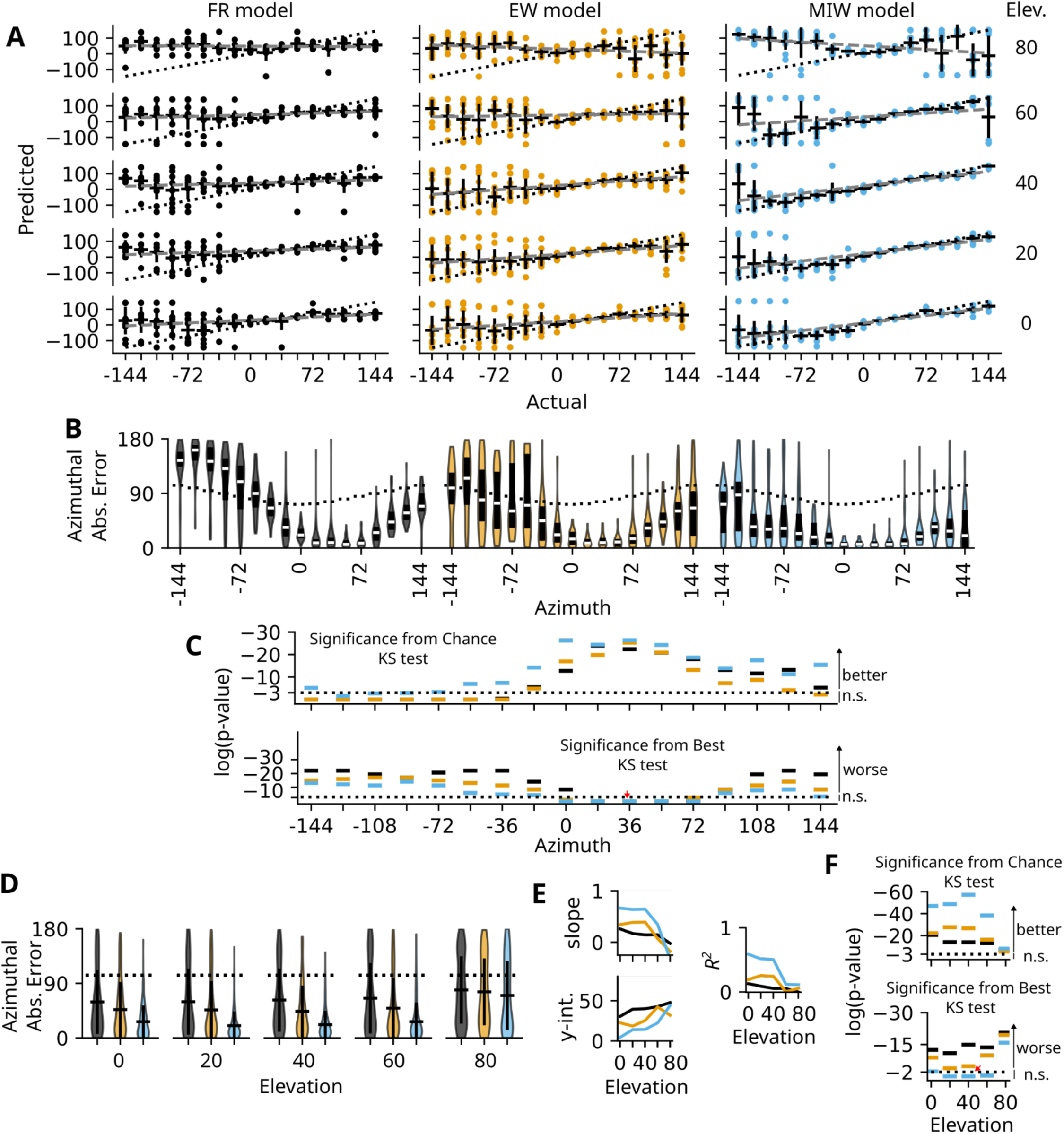
MIW model outperforms FR and EW models on azimuthal error, with best performance in lower elevations in the frontal contralateral auditory space. A) Actual vs Prediction plots for azimuthal results for each model type: FR (black), EW (yellow), and MIW (blue). The Unity line is plotted as a black dotted line. Linear fit lines are shown in dashed grey lines, parameters plotted in D. B) Violin plot for azimuth error based on projected through each azimuth. C) Scatter plot for model performance p-values across the azimuth showing which models show better than random prediction based on a KS test (top) or similar to the best prediction rate (position indicated by the red arrow) based on a KS test (bottom). The dashed line indicates significance. D) Violin plot for azimuth error based on projected through each elevation. E) First-order linear fit parameters and R^2^ values are plotted for linear fits for elevation based on elevation position (shown in A) for FR, EW and MIW models. F) Scatter plot of p-values separated by elevation position, showing that all models, at every elevation, are better than chance for azimuthal errors. However, the low elevations (0°-40°) of the MIW model that does as well as the best azimuthal error rate at 40° elevation.

**Supp Figure 4:**
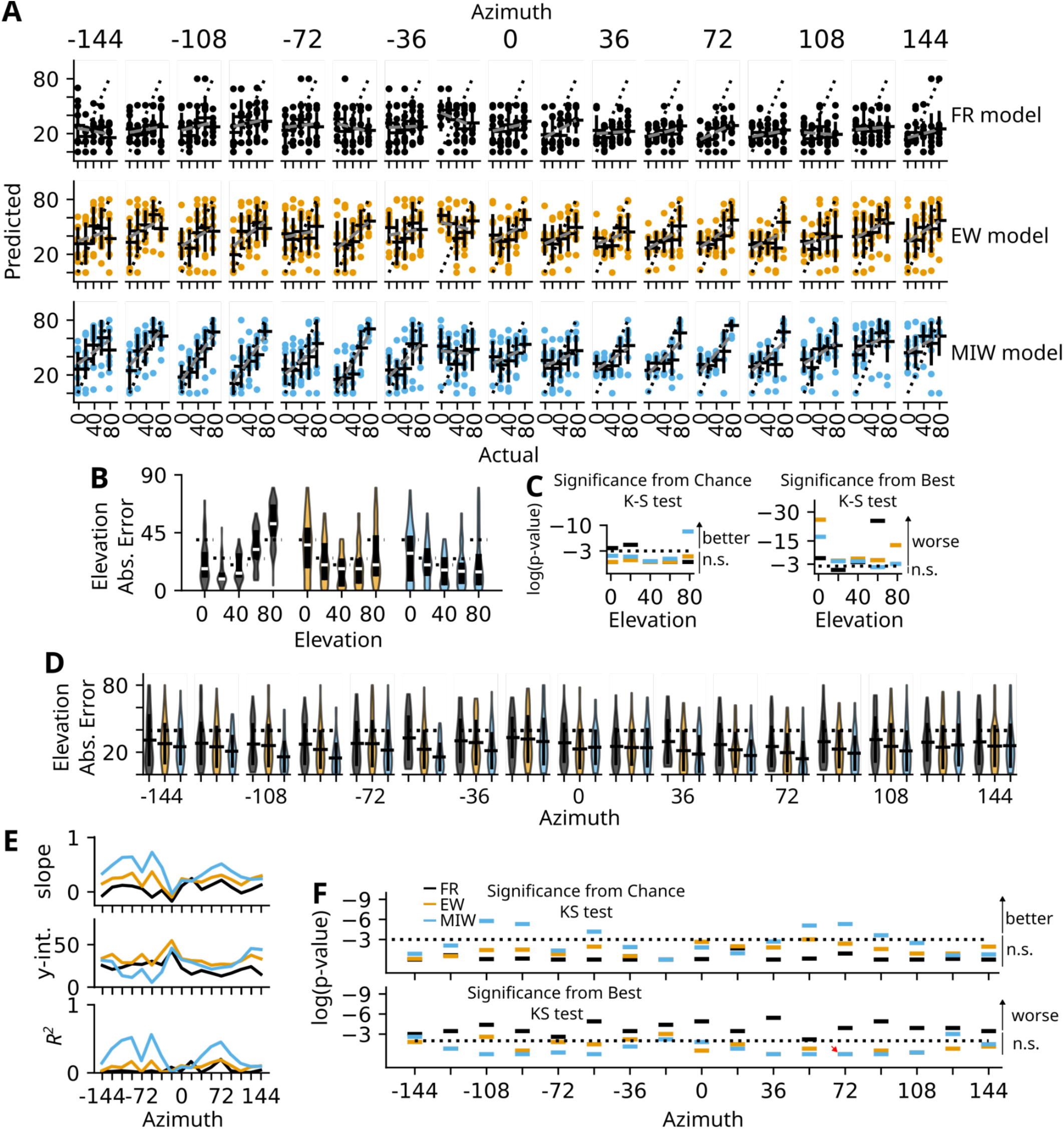
Elevation performance is best at lateral auditory space, with slight improvements in the MI weighted model. A) Actual vs Prediction plots for elevation results for each model type: FR (black), EW (yellow), and MIW (blue). The Unity line is plotted as a black dotted line. Linear fit lines are shown in dashed grey lines, parameters plotted in D. B) Violin plot for elevation error based on projected through each elevation. C) Scatter plot of p-values separated by elevation position, showing which elevation errors are better than randomness or as good as the best azimuthal prediction rate. D) Violin plot for elevation error based on projected through each azimuth. E) First-order linear fit parameters and R^2^ values are plotted for linear fits for elevation based on azimuthal position (shown in A). F) Scatter plot of model performance showing p-values separated by azimuthal position showing which elevation errors are better than random based on a KS test (top) or as good as the best error rate based on a KS test (bottom).

**Supp Figure 5:**
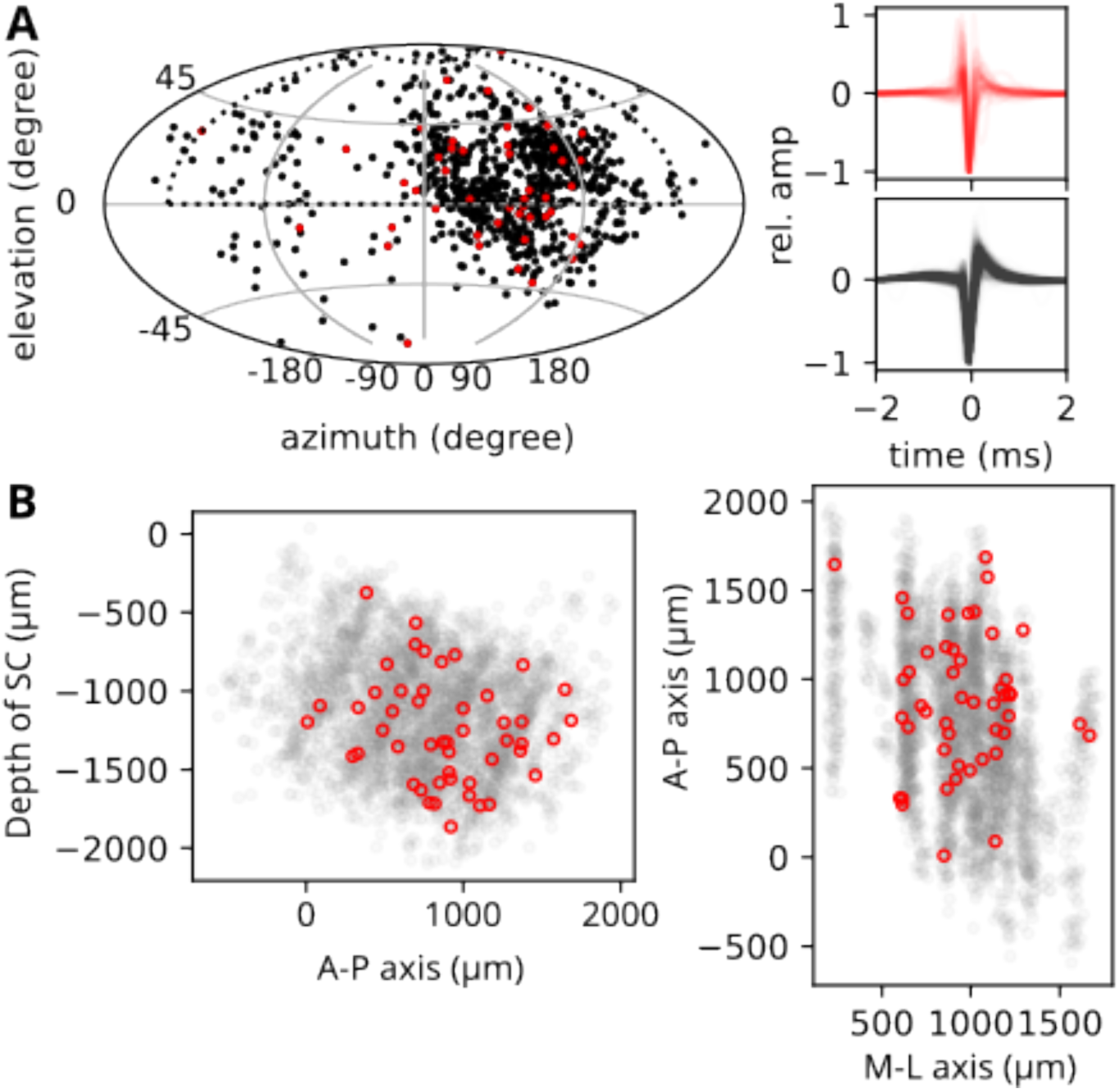
Selection of RF neurons without axonal waveforms. A) All RF neurons (n=813), highlighting the axonal waveforms (n=49; red) and its parameter fit location relative to the remaining RF neurons (n=764; black). Traces of all waveforms are shown (right). B) Anatomical representation of the location of the excluded axonal waveforms on the depth/A-P axis and the M-L/A-P *en face* view showing even dispersion across the SC.

**Supp Figure 6:**
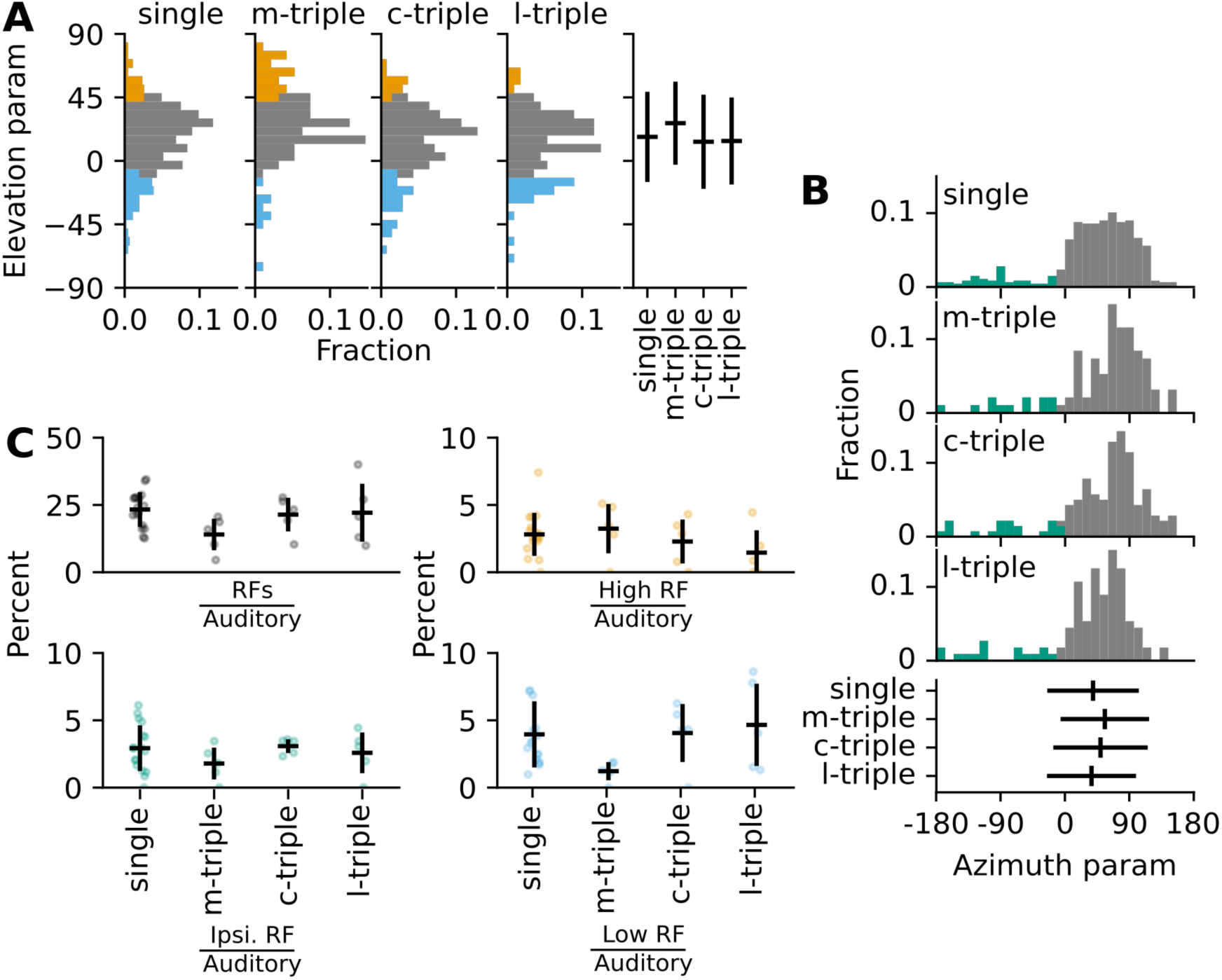
Single insertion experiments have RF neuron characteristics more similar to central and lateral insertion than medial insertion experiments. A) Elevation parameter fits of single insertion into the SC compared to the respective three insertion locations (medial, central, lateral). Five experimental animals were inserted at three locations in the SC (15 total datasets) and the remaining 17 were inserted once. Cutoffs at 45**°** for high and 0° for low elevation RFs. B) Azimuthal parameter fits of single insertion compared to the respective three insertion locations. Cutoff set to -10**°** for ipsilateral RFs as we had many RF on the midline. C) Total number of RF per experimental insertion location (top left), percent counts of neurons that were High RF (top right), Ipsilateral (bottom left), and Low RF neurons (bottom right). Medial insertions tend to have more high and lateral auditory surveilling RF neurons, but fewer neurons.

**SuppTable1:**
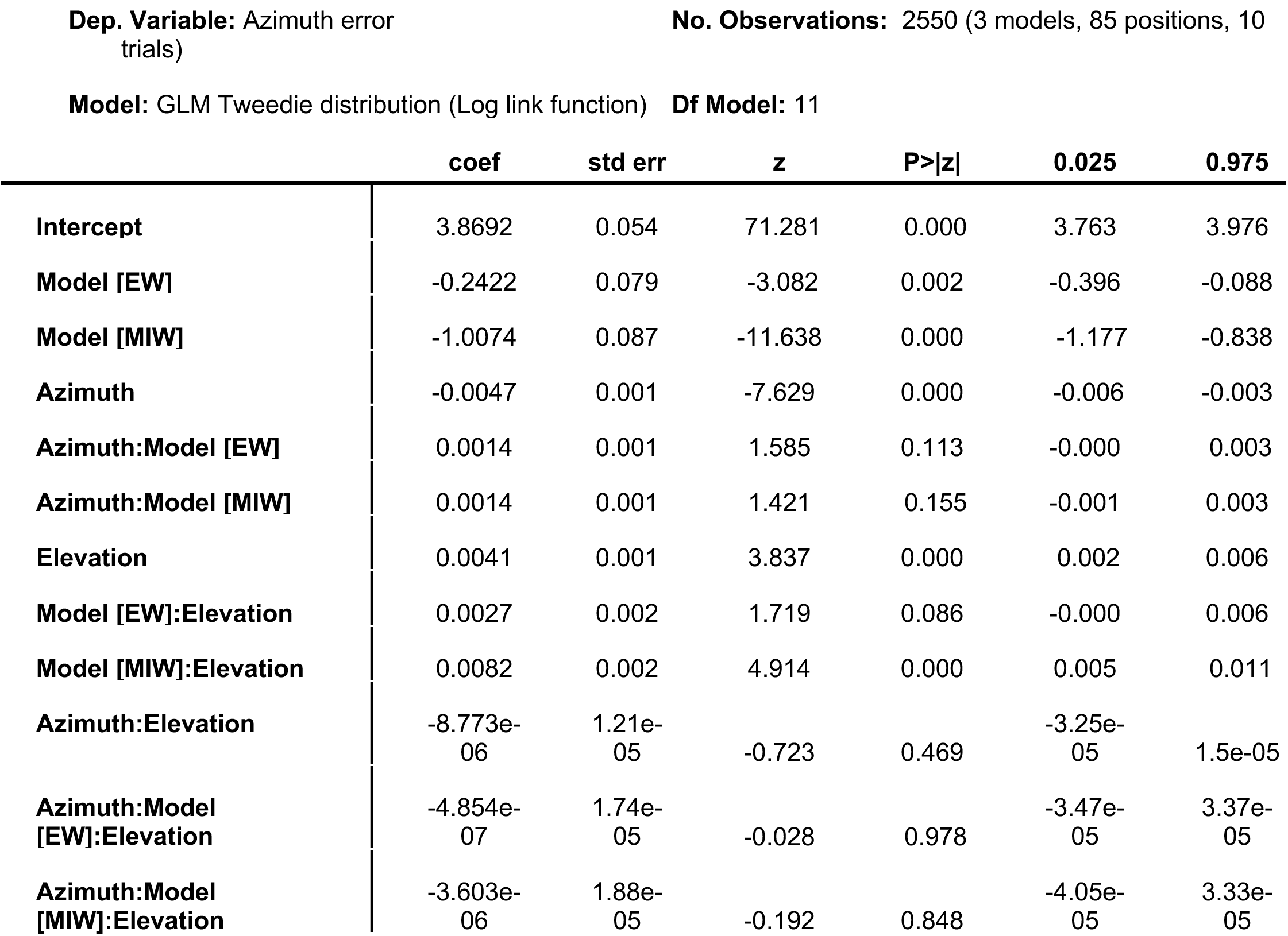
Generalized Linear Model Regression Results for Azimuthal Error.

**SuppTable2:** Generalized Linear Model Regression Results for Elevation Error.

|  | **coef** | **std err** | **z** | **P>\|z\|** | **0.025** | **0.975** |
| --- | --- | --- | --- | --- | --- | --- |
| **Intercept** | 2.7709 | 0.048 | 57.218 | 0.000 | 2.676 | 2.866 |
| **Model [EW]** | 0.5209 | 0.066 | 7.889 | 0.000 | 0.392 | 0.650 |
| **Model [MIW]** | 0.3773 | 0.067 | 5.607 | 0.000 | 0.245 | 0.509 |
| **Azimuth** | -0.0015 | 0.001 | -2.653 | 0.008 | -0.003 | -0.000 |
| **Azimuth:Model [EW]** | 0.0011 | 0.001 | 1.504 | 0.133 | -0.000 | 0.003 |
| **Azimuth:Model [MIW]** | 0.0028 | 0.001 | 3.688 | 0.000 | 0.001 | 0.004 |
| **Elevation** | 0.0128 | 0.001 | 14.055 | 0.000 | 0.011 | 0.015 |
| **Model [EW]:Elevation** | -0.0155 | 0.001 | -11.895 | 0.000 | -0.018 | -0.013 |
| **Model [MIW]:Elevation** | -0.0167 | 0.001 | -12.507 | 0.000 | -0.019 | -0.014 |
| **Azimuth:Elevation** | 2.775e-05 | 1.03e-05 | 2.684 | 0.007 | 7.49e-06 | 4.8e-05 |
| **Azimuth:Model**<br>**[EW]:Elevation** | -<br>3.264e-05 | 1.48e-05 | -2.206 | 0.027 | -6.16e-05 | -3.64e-06 |
| **Azimuth:Model**<br>**[MIW]:Elevation** | -6.101e-05 | 1.52e-05 | -4.022 | 0.000 | -9.07e-05 | -3.13e-05 |

